# cDNA library screening to identify interacting proteins of Golgi-localized type II membrane proteins

**DOI:** 10.1101/2020.02.03.932111

**Authors:** Christian Have Lund, Jennifer R. Bromley, Rosa Laura López-Marqués, Yumiko Sakuragi

## Abstract

In eukaryotes, biosynthesis of many extracellular matrix glycans occurs in the Golgi apparatus. These proteins include glycosyltransferases, modifying enzymes and nucleotide-sugar conversion enzymes, many of which possess a type II membrane topology. Growing evidence indicates that both the function and Golgi localization of many of these proteins are regulated through protein-protein interactions (PPIs). Given the essential nature and conservation of extracellular matrix polysaccharides, it is likely that PPIs are more prevalent among biosynthetic enzymes. However the identification of PPIs among Golgi proteins has been technically challenging due to the generally low abundance and unique membrane topology of these proteins. The aim of this article is to explore the feasibility of cDNA library screening by a yeast-based modified split ubiquitin system as a mean for unbiased screening for PPIs involving a Golgi-localized type II membrane. As a test case, a galacturonosyltransferase1 (GAUT1), involved in pectin biosynthesis in the higher plant Arabidopsis thaliana, was used as the bait. Construction and screening of Arabidopsis cDNA libraries using GAUT1 as the bait successfully led to identification of GAUT7, a previously reported interaction partner of GAUT1, which validates the method. Furthermore, 25 novel candidate interaction partners were identified. The results contribute to shape a field guide for identifying PPIs involving glycan biosynthetic enzymes in eukaryotic cells.

## Introduction

Glycans are important building blocks of life. Glycosylation governs many biological functions in animals, e.g. cell-cell interactions, host-pathogen interactions, inflammation and cancer metastasis. Misregulation of glycan synthesis results in various human genetic diseases [1–3]. In plants, the cell wall is composed of a large and complex extracellular glycan network controlling plant morphology, intercellular communication, and defence against pests amongst other roles [4]. The biosynthesis of many extracellular glycans occurs in the Golgi apparatus. Glycosyltransferases, epimerases, mutases, acetyltransferases, methyltransferases, nucleoside-diphosphatases and nucleotide sugar transporters are among the essential enzymes involved in glycan biosynthesis and are localized to this organelle. Many of the transferases possess a type II membrane topology wherein the membrane anchor domain is flanked by an N-terminal tail localized to the cytosol, and a C-terminal catalytic domain localized to the Golgi lumen [5].

Studies from animals have provided accumulating evidence that PPI is an important principle in spatial and functional regulation of glycan biosynthesis. It has been shown that the glycosyltransferase GlcNAcT-I, involved in N-glycan biosynthesis, and the mannosidase MannII form a high-molecular weight protein complex [6,7]. In this case, complex formation is suggested as a mechanism for targeting of these proteins to the medial sub-compartment of the Golgi apparatus [8,9]. A glycosyltransferase-like protein has been found to act as an inhibitor of GlcNAcT-I by mislocalizing GlcNAcT-I to an earlier Golgi sub-compartment upon association [10]. In heparin biosynthesis, two glycosyltransferases, EXT1 and EXT2, have been shown to form a heterooligomeric complex. Physical association is essential for co-migration of these proteins from the endoplasmic reticulum (ER) to the Golgi apparatus and for enhanced enzymatic activities [11]. Uronosyl-5-epimerase and uronosyl-2-O-sulfatetransferase are also shown to form a complex, which is essential for their localization in the Golgi apparatus [12]. In ganglioside biosynthesis, the glycosyltransferases GalNAcT and GatT2, and GatT1, SialT1 and SialT2 form distinct heterooligomeric complexes [13,14]. Complex formation was found to improve the efficiency of monosialotetrahexosylganglioside synthesis from an exogenously added substrate, suggesting a channelling of the intermediates within the complex [13,14]. Regulatory proteins are also found to interact with glycosyltransferases. Cosmc, a molecular chaperon, interacts with the β3-galactosyltransferase essential for biosynthesis of core 1 O-glycans in animal cell proteoglycans and directly promotes its folding and activity [15,16]. More PPIs are expected to occur between biosynthetic enzymes and regulatory proteins including kinases and proteases [17–19].

In plants, PPIs between cell wall biosynthetic enzymes have been established in cellulose biosynthesis [20–22], in pectic arabinan biosynthesis [23], in glucuronoarabinoxylan biosynthesis [24], and in xyloglucan biosynthesis [25,26]. Among this, a glycosyltransferase, galacturonosyltransferase1 (GAUT1), involved in the pectin homogalacturonan biosynthesis and its homolog GAUT7 interact physically in plant cells via disulphide and non-covalent bonds [27]. This physical interaction plays a pivotal role in targeting GAUT1 to the Golgi apparatus since the N-terminus of GAUT1 is proteolytically processed, releasing the C-terminal catalytic domain from its N-terminal membrane anchor domain. Through interaction with GAUT7, GAUT1 is tethered in the Golgi lumen and thus exerts its biosynthetic activity in synthesizing homogalacturonan [27]. GAUT1 is known to be essential for plant growth and development since homozygous gaut1 mutants have been unobtainable to date [28]. In contrast to the supposedly lethal *gaut1* mutation, the phenotype of the *gaut7* homozygous mutant is not discernible from the wild type [28]. GAUT7 appears to have no enzymatic activity, due to a mutation in the proposed active site [29], which suggests that it may function as a structural protein by anchoring GAUT1 in the Golgi apparatus. Since absence of GAUT1 results in an unviable plant and the loss of GAUT7 results in no severe phenotype, this suggests that further, as yet unidentified proteins act to anchor and retain GAUT1 in the Golgi apparatus.

Identification of PPIs among Golgi-membrane-anchored glycosyltransferases is challenging. Glycosyltransferases generally have low endogenous expression levels and their localization and integration to the membranes of the Golgi apparatus makes detection and heterologous expression often problematic. In addition, each PPI detection method has its limitation. Co-immunoprecipitation requires solubilisation of the membranes with detergents, which potentially interferes with the interaction. Reporter-based interaction assays have been developed. Biomolecular fluorescence complementation can give rise to superb signal-to-noise ration but it is also known to have high rates of false positives due to the irreversible interaction between the reporters [30,31], while Förster resonance energy transfer (FRET) has large technical requirements and low signal-to-noise ratio [32].

The split-ubiquitin system [33] relies on the reconstitution of Cub (C-terminal ubiquitin fragment) and NubG (N-terminal ubiquitin fragment which includes an I13G mutation that decreases the affinity between NubG and Cub) upon interaction of the bait (Cub fused) and prey (NubG fused) proteins. The assembly of the ubiquitin fragments is recognized by cytosolic ubiquitin specific proteases that cleave the C-terminal end of Cub, releasing a synthetic transcription factor (TF). When released, the TF moves into the nucleus and initiates transcription of the reporter genes, HIS3, ADE2 and LacZ. HIS3 and ADE2 encode autotroph selection markers enabling the yeast transformed with proteins engaged in a PPI to grow in media lacking histidine and adenine, while LacZ encodes β-galactosidase that is able to hydrolyse 5-bromo-4-chloro-b-D-galactopyranoside (X-Gal). Both growth on histidine- and adenine-lacking media and blue colour formation in an X-Gal assay is an indication of PPI. The original split-ubiquitin system only allows C-terminal tagging of Cub to the proteins in question [34]. With a type II membrane topology, a common topology of glycosyltransferases located in the Golgi apparatus, C-terminal fusion tags would localize inside the Golgi lumen away from cytosolic ubiquitin specific proteases, thus preventing release of the TF to the nucleus and activation of reporter genes. On the other hand, a modified split-ubiquitin system (Dualsystems Biotech AG Switzerland) allows N-terminal tagging of both Cub and NubG to proteins and can therefore be used to test type II membrane proteins. Indeed, this system has been applied to detect pairwise PPIs amongst cellulose biosynthetic enzymes and KORRIGAN1 (KOR1) [20–22] and between the Golgi-localized type II membrane proteins involved in xyloglucan biosynthesis [25].

Another feature of this modified split-ubiquitin system is its ability to perform cDNA library screens. This allows detection in a high throughput manner of novel PPIs by co-transforming a given bait with a prey cDNA library containing NubG-fused cDNAs prepared from specific tissues, or from combined tissues of a donor organism [35,36]. The cDNA library therefore reflects the transcriptome of the organism from the time and place the mRNA originated. To date, cDNA library screening of interacting partners to Golgi-localized type II membrane proteins has not been reported. This article describes the evaluation of the feasibility of this approach by using GAUT1 as the test case.

## Results and discussion

Firstly, Arabidopsis N-terminal NubG cDNA libraries were screened against full-length GAUT1 N-terminally tagged with a TF-Cub fusion. Prior to screening, GAUT1 was inserted into the bait vector (pBT3-N) and co-expressed in yeast with either Ost1p-NubI, containing a non-mutated form of Nub in fusion with the endomembrane-localized yeast protein Ost1p [37], or cytosolic NubG (empty prey vector, pPR3-N) to test for functionality and auto-activation. According to this test TF-Cub-GAUT1 is functional as bait, which can be inferred from both growth and blue colour formation when co-transformed with Ost1p-NubI (Figure 1a). In addition, no reporter growth and blue colour formation was observed when the GAUT1 fusion was co-expressed with the cytosolic NubG expressed from the empty prey vector (pPR3-N), indicating no detectable auto-activation (Figure 1a).

**Figure 1.**
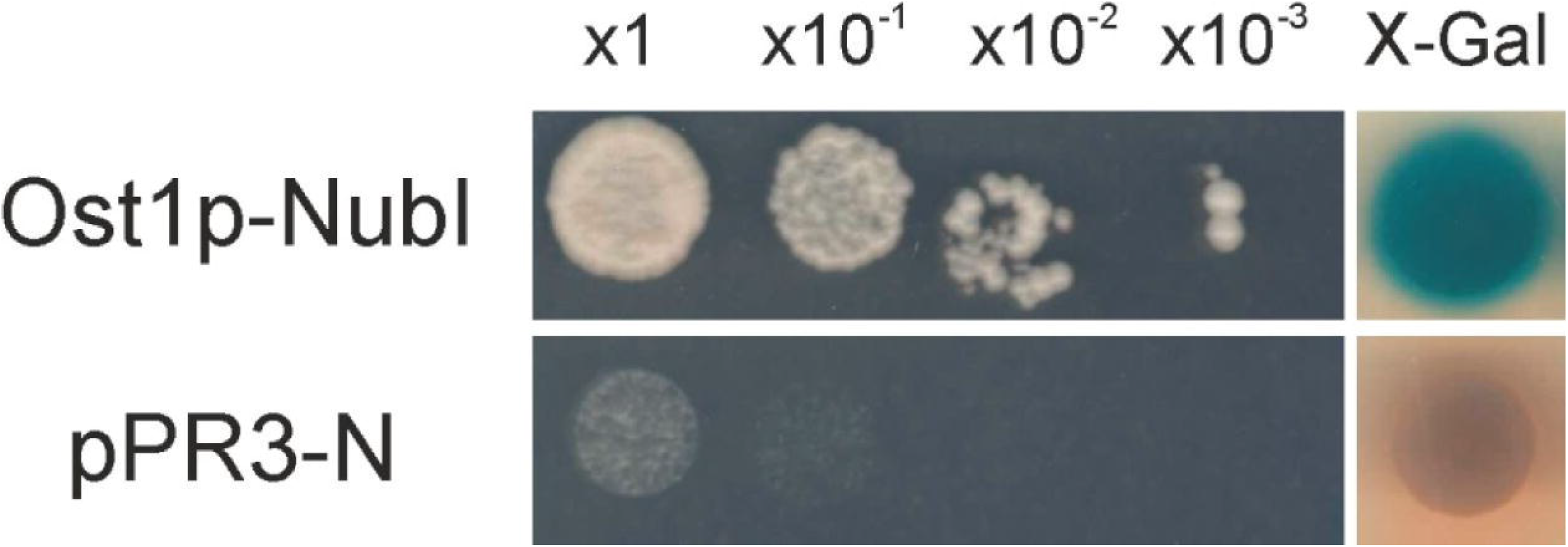
Functionality test of TF-Cub-GAUT1. The yeast strain NMY51 bearing TF-Cub-GAUT1 and Ost1p-NubI or pRR3-N were growth on the solid SD-L-W-H-A medium after serial dilutions (left) and the SD-L-W medium containing X-Gal (right).

Three cDNA libraries were generated containing the transcriptome of Arabidopsis leaves, stems, and flowers. Yeast were first transformed with the plasmid bearing TF-Cub-GAUT1 and subsequently transformed with the various cDNA libraries, giving rise to >2×10^6^ independent double transformants on a SD-L-W medium (synthetic medium lacking leucine and histidine, allowing selection of transformants bearing both the prey and the bait), sufficient for covering the entire Arabidopsis transcriptome according to manufacturer′s instructions. After selection for PPIs on a SD-L-W-H-A medium (a SD-L-W medium lacking histidine, and adenine), 180 yeast colonies were recovered. The gene products responsible for facilitating the interactions were identified by sequencing of the cDNA followed by analysis using the Basic Local Alignment Search Tool (BLAST) [38]. The distribution of their functional categories is summarised in Table 1. An extended list of identified interactors is provided in Table S1.

**Table 1.** Overview of yeast colonies picked, interactions identified and their involvement in relation to GAUT1.

|  |  |  |
| --- | --- | --- |
| Growing yeast colonies picked |  | 180 |
| Number of non-redundant proteins identified |  | 52 |
| Number of non-redundant proteins localized to nucleus, chloroplasts, and mitochondria |  | 27 |
| Candidate interacting proteins |  | 25 |
| By category | Cell wall biosynthesis | 4 |
|  | Vesicle transport | 7 |
|  | Other function | 8 |
|  | Unknown function | 6 |

Notably GAUT7 was identified. This result was expected given the reported PPI between GAUT1 and GAUT7 [27]. Then, full-length GAUT7 was fused with NubG and PPI between TF-Cub-GAUT1 and NubG-GAUT7 was tested. Both growth on SD-L-W-H-A medium and blue colour development after X-Gal assay were detected (Figure 2a). It is unlikely that NubG-GAUT7 is randomly interacting with Golgi-localizing proteins, as we could not detect any PPI with Cub-fused Anp1p, a yeast Golgi-localized enzyme involved in N-glycan biosynthesis [25] (Figure 2b). These results show that cDNA library screening using the modified split ubiquitin system can successfully identify a known PPI and therefore validate the method.

**Figure 2.**
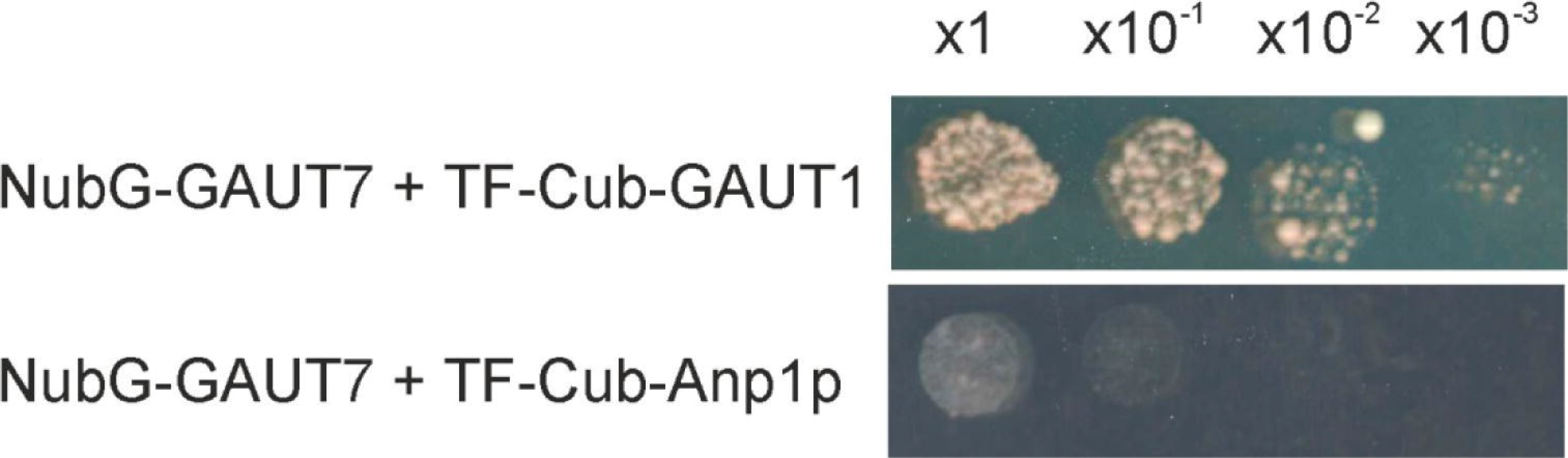
NubG-GAUT7 interacts specifically with TF-Cub-GAUT1. The yeast strain NMY51 bearing NubG-GAUT7 and TF-Cub-GAUT1 or TF-Cub-Anp1p were grown on the solid SD-L-W-H-A medium after serial dilutions. X-Gal assay was omitted due to occasional leakiness of the reporter.

The identities of 52 genes were determined out of the 180 yeast colonies recovered. Twenty-seven of them were considered as potential false positives due to their known or predicted localization to nucleus, chloroplast, peroxisome, and mitochondria (Table S1). Most of these proteins were involved in processes such as energy metabolism, DNA modulation and regulatory pathways. The remaining 25 gene products were considered candidate GAUT1 interactors. These putative interactors could be sorted according to their Gene Ontology (GO) terminology into four groups: cell wall biosynthesis, vesicle transport, other function, and unknown function (Table 1). It is important to note that validation of PPIs between GAUT1 and these interactors is pending; however, as discussed below, an initial look into the list may provide insights into the cell biology and possible coordination of glycan biosynthesis in the Golgi apparatus.

The xyloglucan:xylosyltransferase1 (XXT1) was identified twice in this screen to interact with GAUT1. XXT1 is involved in the biosynthesis of the hemicellulose xyloglucan and is also located in the Golgi apparatus [39,40]. Notably, it has been previously shown that xyloglucan is covalently linked to pectins via the reducing end of the arabinogalactan sidechain of rhamnogalacturonan I in a wide range of plant species [41–46]. Consistently, Arabidopsis mutants defective in xyloglucan biosynthesis (*xxt1* and *xxt2* single and double mutants) were found to contain significantly decreased levels of rhamnose and arabinose, the monosaccharides that constitute rhamnogalacturonan I [47]. It appears that the linkage is formed during biosynthesis of pectin and xyloglucan inside the Golgi lumen [45]. Identification of XXT1 in the screen might suggest that homogalacturonan and xylogalacturon biosynthesis are also be spatially organized in the Golgi apparatus and have an impact on the formation of the pectin-xyloglucan cross-link.

Another interesting protein identified in the screen was CHITINASE LIKE (CTL) 2. This protein and CTL1, a functionally equivalent homolog of CTL2, have been shown to bind cellulose and hemicelluloses and impact the formation of cellulose microfibrils [48]. Furthermore, CTL1 was shown to co-localize with the cellulose synthase in the Golgi apparatus and in post-Golgi compartments called MASCs/SmaCCs [48], and the *ctl1* mutant is known to cause microtubule disorganization [49]. It is noteworthy that a similar defect in microtubule disorganization has been reported for a mutant of KORRIGAN1 that has been shown to form complexes with a cellulose synthase [22] and a homogalacturonan synthase involving GAUT1 [27]. These observations suggest that biosynthesis and/or assembly of pectins, hemicelluloses, and cellulose may be coordinated in the endomembrane system through a yet unknown mechanism that may involve the above-mentioned proteins.

Several proteins involved in intracellular vesicle transport were also identified in the library screen. Vesicle transport is required to for the proper targeting of glycan biosynthetic enzymes from the ER to the Golgi apparatus via the anterograde transport and within the Golgi apparatus or from the Golgi apparatus to the ER via the retrograde transport [50–52]. Furthermore, the newly synthesized glycan polymers need to be transported to the extracellular matrix. An ER lumen protein-retaining receptor family protein was identified twice in the screen. This protein might be important in sorting properly folded GAUT1 to be engulfed into a forming vesicle. ADP-ribosylation factor A1E (ARFA1E) and secretory carrier membrane protein (SCAMP) were also identified and may be involved in recognition of GAUT1 during vesicle formation, while Profilin-2 (PFN2) facilitates transport of the vesicle to the cis cisternae of the Golgi via actin filaments.

LPAT2 is a lysophosphatidyl acyltransferase involved in synthesis of phosphatidic acids that constitute phospholipids in cell membranes [53]. In the domain synthesis model of pectin [5], oligo- or polysaccharides primers are synthesized from sugar nucleotides or potentially from lipid-linked sugars and elongated before being transferred *en bloc* to form polysaccharides. It has been shown that phosphoinositides that are synthesized from phosphatidic acids exert a significant impact on the secretion of pectins [54]. It might be that LPAT2 plays a role in pectin secretion through complex formation with GAUT1 and further studies are required to test this possibility.

## Summary and conclusions

cDNA library screening can facilitate unbiased identification of PPIs and here we show that the method can be applied for Golgi-localized type II membrane proteins. Needless to say that most methods are not perfect and the split-ubiquitin system is not an exception. It appears that a high number of false positive interaction partners can be identified [55], which also is the case in our library screens. Twenty-seven of the observed interactors were considered as false positives due to their subcellular localization (Table S1). Notably, eight of the chloroplastic proteins identified here were also found in another screen using a bait localizing to the chloroplast [56]. These can be detected by random interaction tests or by in silico analysis, using tools such as BioGRID [57], and eliminated from an interaction network. We have also previously observed a high number of false negatives among PPIs involving Golgi-localizing type II membrane proteins [25]. In spite of these caveats, this system can quickly provide a list of potentially interacting candidates, which is an excellent starting point to identify an interaction network. Pectin biosynthesis takes place inside the Golgi lumen and pectin polymers are then transported from the Golgi stacks to the apoplast where they are integrated with the other cell wall polymers. Identification of PPIs can set the ground for further investigation of the mechanism of glycan biosynthesis and assembly in eukaryotic cells.

## Materials and Methods

### Construction and functionality test of TF-Cub-GAUT1

The modified split-ubiquitin assay was performed using *Saccharomyces cerevisiae* yeast strain NMY51 (MAT*a his3Δ200 trp1–901 leu2–3,112 ade2* LYS2::(lexAop)4-HIS3 *ura3*::(lexAop)8-lacZ *ade2*::(lexAop)8-ADE2 GAL) [58] using pBT3-N (bait), Ost1p–NubI, and pPR3-N (prey) vectors (Dualsystems Biotech AG, Schlieren, Switzerland) to perform the functionality test.

For cloning GAUT1 into pBT3-N, the corresponding coding sequence was amplified by PCR using the following oligonucleotide primers: 5’-GGCCATTACGGCCATGGCGCTAAAGCGAGGGC-3’ and 5’-GGCCGAGGCGGCCTTATTCATGAAGGTTGCAACG-3’, where the underlined sequence contains asymmetric SfiI recognition sites to allow directional cloning. Both the PCR product and the pBT3-N vector were digested wtih *SfiI* and ligated with T4 DNA ligase. The resulting plasmid was designated as pTF-Cub-GAUT1.

The vectors were introduced in pairs into NMY51 by LiAc transformation [59]. Transformants were selected on Synthetic Defined (SD) media plates (0.7% Yeast Nitrogen base, 2% glucose, 1.5% agar), containing yeast drop out supplements lacking leucine and, tryptophan) (SD-L-W).Strains carrying both vectors were grown in liquid SD-L-W (same as above without agar) to OD546nm of 1.5. Serial dilutions (from 1 to 1000 folds) were spotted on SD-L-W, SD-L-W-H (lacking leucine, tryptophan and histidine), and SD-L-W-H-A (lacking leucine, tryptophan, histidine and adenine). Growth on the SD-L-W-H and SD-L-W-H-A plates was interpreted as an indication of interaction. Yeast cells growing on SD-L-W plates were tested for β-galactosidase activity using the X-gal overlay assay [25].

### Generation of cDNA libraries

Total RNA was extracted from leaves, stems and flowers of old 6-week old *Arabidopsis thaliana* (ecotype Columbia-0) plants grown under a long day photoperiod (16 h light/8 h dark, 60% humidity) using the SpectrumTM Total RNA extraction kit (Sigma-Aldrich, USA). The sample was enriched for mRNA using the Poly(A) Purist MAG kit (Invitrogen, USA). The cDNA libraries were generated and normalized using the EasyClone normalized cDNA library construction kit (Dualsystems Biotech, Switzerland) according to manufacturer′s instructions. Finally normalized double-stranded cDNA and the pPR3-N vector were digested with SfiI and subsequently ligated with T4 DNA ligase to generate the N-terminally NubG fused prey cDNA libraries. The cDNA libraries were amplified in DH10B E.coli cells and purified using QIAGEN Plasmid Mega Purification kit (Qiagen, Germany). The average size of the cDNA inserts into pPR3-N was evaluated by purifying 20 random colonies each and digesting with *SfiI*, and ranged from approximately 0.9kb to 1.8kb.

### cDNA library screen

pTF-Cub-GAUT1 was introduced into NMY51 as previously described [59] and selected on SD-L plates at 30°C for 3 days. The introduction of the cDNA libraries into NMY51 carrying pTF-Cub-GAUT1 was performed as described in Library transformation and selection of interactors in DUALmembrane starter kit (Dualsystems Biotech, Switzerland) and selected on SD-L-W-H-A plates. Colonies were picked and grown in 5 ml liquid SD-L-W medium and the prey vectors were isolated using the GenElute Plasmid Miniprep Kit (Sigma-Aldrich, USA) after cell wall disruption. For this, yeast cells in resuspension buffer were mixed with 0.3 g acid-washed glass beads and shaken in a Retch mixer mill for 4 min, before addition of lysis buffer.

The recovered prey plasmids were further amplified in *E. coli* and isolated as described above and inserts were identified by direct DNA sequencing. Alternatively, the inserts were amplified by PCR from the recovered prey plasmids using the following oligonucleotide primers: NubG forward sequencing primer, 5’-TGGAAGTTGAATCTTCCGATAC-3’; and DUAL reverse sequencing primer, 5’-AAGCGTGACATAACTAATTAC-3’. PCR products were gel extracted using the freeze-squeeze DNA gel extraction method [60,61], before sequencing. The obtained DNA sequences were searched against the non-redundant nucleotide database using NCBI Basic Local Alignment Search Tool (BLAST) [38].

## Supplementary material

Table S1. Identified interaction partners to GAUT1 in the modified split-ubiquitin cDNA library screen

## Acknowledgments

This work was supported by the Danish Advanced Technology Foundation [Biomass for the 21st century, grant number 001-2011-4], the Danish Council for Strategic Research [Plant Power, grant number 12-131834], the Danish Council for Independent Research | Natural Sciences [FNU, project number 10-083406] and European Union′s Seventh Framework Programme FP7 (ENERGY-2010-1 DirectFuel, grant number 256808). We thank Randi E. Rasmussen, Jonas Gensby, Annemette Lyhne-Kjærbye, Marc Pilegaard, Max Woveries and Simon Erstad for experimental assistance. The authors declare no competing financial interests.

