## Supplemental Table 1 for "cDNA library screening to identify interacting proteins of Golgi-localized type II membrane proteins"

**Table S1. Identified interaction partners to GAUT1 in the modified membrane split-ubiquitin cDNA library screen**

| **AGI** | **Description** | **SUBAcon** | | **Ref** | | **GO terms** | | | | | | |
| --- | --- | --- | --- | --- | --- | --- | --- | --- | --- | --- | --- | --- |
|  |  |  |  |  |  | **Cellular component** | | | **Molecular function** | **Biological process** | | |
| **Cell wall biosynthesis** | | | | | | | | | | | | |
| AT2G38650 | α-1,4-galacturonosyltransferase (GAUT7): Encodes protein involved in carbohydrate biosynthesis and annotated to have polygalacturonate 4-alpha-galacturonosyltransferase activity | | Golgi | | [1,2] | Golgi apparatus (GO:0005794)  Trans-Golgi network (GO:0005802) | Polygalacturonate 4-α-galacturonosyltransferase activity (GO:0047262)  Transferase activity,  transferring glycosyl groups (GO:0016757) | | | | Carbohydrate biosynthetic process (GO:0016051)  Hydrogen peroxide biosynthetic process (GO:0050665)  Protein desumoylation (GO:0016926) | |
| AT3G62720 | Xyloglucan 6-xylosyltransferase (XXT1): Encodes a protein with xylosyltransferase activity in xyloglucan biosynthesis | | Golgi | | [3,4] | Golgi apparatus (GO:0005794)  Trans-Golgi network (GO:0005802)  Integral to membrane (GO:0016021) | Transferase activity, transferring glycosyl groups (GO:0016757)  UDP-xylosyltransferase activity (GO:0035252)  Xyloglucan 6-xylosyltransferase  activity (GO:0033843) | | | | Xyloglucan biosynthetic process (GO:0009969)  Polysaccharide biosynthetic process (GO:0000271)  Response to chitin (GO:0010200)  Root hair elongation (GO:0048767)  Response to mechanical stimulus (GO:0009612) | |
| AT5G11740 | Arabinogalactan protein (AGP15) | | Extracellular (Plasma membrane) | | [5] | Plasma membrane (GO:0005886)  Anchored to membrane (GO:0031225) |  | | | |  | |
| AT3G16920 | CHITINASE-LIKE PROTEIN 2 (CTL2): Encodes a chitinase-like protein with chitinase activity and involved in lignin biosynthesis. | | Extracellular (Plasma membrane) | | [6,7] | Plasma membrane (GO:0005886)  Extracellular region (GO:0005576) | Chitinase activity (GO:0004568) | | | | Xylan biosynthetic process (GO:0045492)  Hydrogen peroxide biosynthetic process (GO:0050665)  Protein desumoylation (GO:0016926)  Glucuronoxylan metabolic process (GO:0010413)  Lignin biosynthetic process (GO:0009809)  Cell wall macromolecule catabolic process (GO:0016998)  Carbohydrate metabolic process (GO:0005975)  Vegetative to reproductivephase transition of meristem (GO:0010228) | |
| **Vesicle transport** | | | | | | | | | | | | |
| AT4G38790 | ER lumen protein retaining receptor family protein involved in protein retention in ER and transport | | ER | |  |  | ER retention sequence binding (GO:0046923)  Receptor activity (GO:0004872) | | | | Protein transport (GO:0015031)  Protein retention in ER lumen (GO:0006621) | |
| AT3G62290 | ADP-ribosylation factor A1E (ARFA1E): A member of ARF GTPase family and is essential for vesicle coating and uncoating and functions in GTP-binding. | | Golgi (ER, cytosol, vacuole, plasma membrane) | |  | Golgi apparatus (GO:0005794)  Vacuole (GO:0005773)  Intracellular (GO:0005622)  Plasmodesma (GO:0009506) | GTP binding (GO:0005525)  Phospholipase activator activity (GO:0016004) | | | | Protein transport (GO:0015031)  Intracellular protein transport (GO:0006886)  Small GTPase mediated signal transduction (GO:0007264)  N-terminal protein myristoylation (GO:0006499) | |
| **AGI** | **Description** | | **SUBAcon** | | **Ref** | **GO terms** | | | | | | |
|  |  |  |  |  |  | **Cellular component** | | **Molecular function** | | | | **Biological process** |
|  |  | |  | |  | **Vesicle transport** **continued** | |  | | | |  |
| AT1G04750† | Vesicle-associated membrane protein 721 (VAMP721): Encodes vesicle-associated membrane protein 7B (VAMP7B, or VAMP721). Required for cell plate formation | | Plasma membrane (Golgi) | |  | Plasma membrane (GO:0005886)  Integral to membrane (GO:0016021)  Endosome (GO:0005768)  Membrane (GO:0016020)  Plasmodesma (GO:0009506) | Molecular_function_unknown (GO:0003674) | | | | Vesicle-mediated transport (GO:0016192)  Protein targeting to plasma membrane (GO:0072661)  Cell plate (GO:0009504)  Cell plate formation involved in planttype cell wall biogenesis (GO:0009920)  Transport (GO:0006810) | |
| AT4G29350 | Profilin 2 (PFN2): Actin monomer-binding protein that regulates the organization of actin cytoskeleton | | Cytosol (ER) | |  | Actin cytoskeleton (GO:0015629)  Chloroplast (GO:0009507) Cytoplasm (GO:0005737)  Plasma membrane (GO:0005886) | Protein binding (GO:0005515)  Actin binding (GO:0003779) | | | | Cytoskeleton organization (GO:0007010)  Actin polymerization or depolymerization (GO:0008154)  Actin cytoskeleton organization (GO:0030036) | |
| AT1G11180 | Secretory carrier membrane protein (SCAMP): Involved in protein transport and has transmembrane transporter activity | | Plasma membrane | |  | Plasma membrane (GO:0005886)  Integral to membrane (GO:0016021) | Transmembrane transporter activity (GO:0022857) | | | | Protein transport (GO:0015031) | |
| AT1G17080† | Ribosomal protein L18ae-like protein: Involved in vesicle-mediated transport to target proteins to vacuole. | | Plasma membrane (vacuole) | |  |  | Protein binding (GO:0005515)  Structural constituent of ribosome (GO:0003735) | | | | Biological_process_unknown (GO:0008150)  Vesicle-mediated transport (GO:0016192)  Protein targeting to vacuole (GO:0006623)  Translation (GO:0006412) | |
| AT5G40780 | Lysine histidine transporter (LHT1): Encodes a high-affinity transporter for cellular amino acid and involved in ER to Golgi vesicle-mediated transport, amino acid transmembrane transport and in response to karrikin | | Plasma membrane | |  | Plasma membrane (GO:0005886)  Membrane (GO:0016020) | Amino acid transmembrane transporter activity (GO:0015171) | | | | Amino acid transport (GO:0006865)  Cellular membrane fusion (GO:0006944)  ER to Golgi vesicle-mediated transport (GO:0006888)  Anion transport (GO:0006820)  Amino acid import (GO:0043090)  N-terminal protein myristoylation (GO:0006499)  Ammonium transport (GO:0015696)  Endoplasmic reticulum unfolded protein response (GO:0030968)  Protein targeting to membrane (GO:0006612)  Response to karrikin (GO:0080167)  Regulation of plant-type hypersensitive response (GO:0010363)  Nucleotide transport (GO:0006862)  Negative regulation of programmed cell death (GO:0043069)  Basic amino acid transport (GO:0015802) | |
| **AGI** | **Description** | | **SUBAcon** | | **Ref** | **GO terms** | | | | | | |
|  |  |  |  |  |  | **Cellular component** | | **Molecular function** | | | | **Biological process** |
| **Others** | | | | | | | | | | | | |
| AT3G57650 | Lysophosphatidyl acyltransferase 2 (LPAT2): Encodes an endoplasmic reticulum localized protein with lysophosphatidyl acyltransferase activity. | | ER | | [8] | Endoplasmic reticulum (GO:0005783)  Extracellular region (GO:0005576) | Transferase activity, transferring acyl groups (GO:0016746)  1-acylglycerol-3-phosphate O-acyltransferase activity (GO:0003841) | | | | Metabolic process (GO:0008152)  Phosphatidylglycerol biosynthetic process (GO:0006655) | |
| AT3G15353 | Metallothionein 3 (MT3): Involved in limitation of oxidative damage by binding and detoxifying excess copper and other metals | | Plasma membrane (ER, Golgi, extracellular, cytosol) | |  | Extracellular region (GO:0005576) | Copper ion binding (GO:0005507) | | | | Pattern specification process (GO:0007389)  Water transport (GO:0006833)  Protein targeting to membrane (GO:0006612)  Cellular copper ion homeostasis (GO:0006878)  Root morphogenesis (GO:0010015)  Auxin polar transport (GO:0009926)  Regulation of cell size (GO:0008361)  Positive regulation of flavonoid biosynthetic process (GO:0009963)  Regulation of planttype hypersensitive response (GO:0010363)  Response to salt stress (GO:0009651)  Response to osmotic stress (GO:0006970)  Response to cadmium ion (GO:0046686)  Response to fructose stimulus (GO:0009750)  Growth (GO:0040007) | |
| AT3G16240 | Delta tonoplast integral protein (DELTA-TIP): Involved in water, ammonium and urea transport | | Vacuole | |  | Golgi apparatus (GO:0005794)  Cytoplasm (GO:0005737)  Plasma membrane (GO:0005886)  Plant-type cell wall (GO:0009505)  Cell wall (GO:0005618)  Membrane (GO:0016020)  Plasmodesma (GO:0009506)  Vacuolar membrane (GO:0005774)  Central vacuole (GO:0042807)  Vacuole (GO:0005773)  Protein storage vacuole (GO:0000326)  Chloroplast (GO:0009507)  Chloroplast envelope (GO:0009941) | Methylammonium transmembrane transporter activity (GO:0015200)  Transporter activity (GO:0005215)  Water channel activity (GO:0015250)  Urea transmembrane transporter activity (GO:0015204)  Ammonia transmembrane transporter activity (GO:0051739) | | | | Divalent metal ion transport (GO:0070838)  Cellular cation homeostasis (GO:0030003)  Water transport (GO:0006833)  Urea transport (GO:0015840)  Response to salt stress (GO:0009651)  Transport (GO:0006810)  Calcium ion transport (GO:0006816)  Golgi organization (GO:0007030)  Trans-membrane transport (GO:0055085) | |
| AT2G30860 | Glutathione S-transferase (GSTF9): Encodes glutathione transferase belonging to the phi class of GSTs | | Cytosol | |  | Plasma membrane (GO:0005886)  Cytoplasm (GO:0005737)  Cytosol (GO:0005829)  Plasmodesma (GO:0009506)  Thylakoid (GO:0009579)  Chloroplast stroma (GO:0009570)  Chloroplast (GO:0009507)  Apoplast (GO:0048046)  Vacuole (GO:0005773) | Glutathione peroxidase activity (GO:0004602)  Glutathione transferase activity (GO:0004364)  Glutathione binding (GO:0043295)  Copper ion binding (GO:0005507) | | | | Systemic acquired resistance (GO:0009627)  Regulation of defense response (GO:0031347)  Indoleacetic acid biosynthetic process (GO:0009684)  Defense response to bacterium (GO:0042742)  Response to cadmium ion (GO:0046686)  Toxin catabolic process (GO:0009407)  Tryptophan catabolic process (GO:0006569)  Glycine catabolic process (GO:0006546)  Defense response (GO:0006952)  Response to zinc ion (GO:0010043)  Response to osmotic stress (GO:0006970) | |
| **AGI** | **Description** | | **SUBAcon** | | **Ref** | **GO terms** | | | | | | |
|  |  |  |  |  |  | **Cellular component** | | **Molecular function** | | | | **Biological process** |
| **Others continued** | | | | | | | | | | | | |
| AT1G65980 | THIOREDOXIN-DEPENDENT PEROXIDASE 1 (TPX1): Protein with antioxidant activity | | Cytosol | |  | Membrane (GO:0016020)  Cytoplasm (GO:0005737)  Chloroplast (GO:0009507)  Cytosol (GO:0005829)  Plasma membrane (GO:0005886) | Oxidoreductase activity (GO:0016491) | | | | Toxin catabolic process (GO:0009407)  Response to cadmium ion (GO:0046686)  Antioxidant activity (GO:0016209) | |
| AT1G59870 | ABC transporter G family member 36 (PEN3): ATP binding cassette transporter. | | Plasma membrane | |  | Plasma membrane (GO:0005886)  Membrane (GO:0016020)  Mitochondrion (GO:0005739)  Chloroplast (GO:0009507)  Chloroplast envelope (GO:0009941)  Vacuolar membrane (GO:0005774) | ATP binding (GO:0005524)  Cadmium ion transmembrane transporter activity (GO:0015086)  Nucleotide binding (GO:0000166)  ATPase activity, coupled to transmembrane movement of substances (GO:0042626)  Nucleoside-triphosphatase activity (GO:0017111)  ATPase activity (GO:0016887) | | | | Defense response to bacterium (GO:0042742)  Regulation of multiorganism process (GO:0043900)  Regulation of response to biotic stimulus (GO:0002831)  Divalent metal ion transport (GO:0070838)  Cadmium ion transport (GO:0015691)  Detection of biotic stimulus (GO:0009595)  Negative regulation of defense response (GO:0031348)  Cellular response to nitrogen starvation (GO:0006995)  Cellular response to indolebutyric acid stimulus (GO:0071366)  Regulation of defense response (GO:0031347)  Drug transmembrane transport (GO:0006855)  Salicylic acid biosynthetic process (GO:0009697)  Indole glucosinolate catabolic process (GO:0042344)  Systemic acquired resistance (GO:0009627)  Response to chitin (GO:0010200)  Defense response to fungus, incompatible interaction (GO:0009817)  Response to abscisic acid stimulus (GO:0009737)  Defense response to fungus (GO:0050832)  Regulation of hydrogen peroxide metabolic process (GO:0010310)  Protein targeting to membrane (GO:0006612)  MAPK cascade (GO:0000165)  Cellular cation homeostasis (GO:0030003)  Jasmonic acid mediated signaling pathway (GO:0009867)  Regulation of plant-type hypersensitive response (GO:0010363)  Systemic acquired resistance, salicylic acid mediated signaling pathway (GO:0009862)  Defense response by callose deposition in cell wall (GO:0052544)  Positive regulation of flavonoid biosynthetic process (GO:0009963) | |
| **AGI** | **Description** | | **SUBAcon** | | **Ref** | **GO terms** | | | | | | |
|  |  |  |  |  |  | **Cellular component** | | **Molecular function** | | | | **Biological process** |
| **Others continued** | | | | | | | | | | | | |
| AT5G39740 | 60S ribosomal protein L5-2 (RPL5B): Involved in ribosome biogenesis and plays a role in organ size control by promoting cell proliferation and preventing compensation in normal leaf development | | Cytosol | |  | Cytosolic ribosome (GO:0022626)  Intracellular (GO:0005622)  Cytoplasm (GO:0005737)  Ribosome (GO:0005840)  Nucleolus (GO:0005730)  Vacuole (GO:0005773)  Cytosol (GO:0005829)  Sytosolic large ribosomal subunit (GO:0022625)  Plasma membrane (GO:0005886) | 5S rRNA binding (GO:0008097)  Structural constituent of ribosome (GO:0003735) | | | | Glycolysis (GO:0006096)  Translation (GO:0006412)  Response to cadmium ion (GO:0046686)  Cell proliferation (GO:0008283)  RNA methylation (GO:0001510)  Ribosome biogenesis (GO:0042254)  Gluconeogenesis (GO:0006094)  Leaf morphogenesis (GO:0009965)  Response to salt stress (GO:0009651)  Pyrimidine ribonucleotide biosynthetic process (GO:0009220) | |
| AT3G18280† | Bifunctional inhibitor/lipid-transfer protein/seed storage 2S albumin superfamily protein involved in lipid binding and transport | | Extracellular | |  | Extracellular region (GO:0005576) |  | | | | Regulation of plant-type hypersensitive response (GO:0010363)  Lipid transport (GO:0006869)  Positive regulation of flavonoid biosynthetic process (GO:0009963)  Protein targeting to membrane (GO:0006612) | |
| **Unknown function** | | | | | | | | | | | | |
| AT5G43460† | HR-like lesion-inducing protein with unknown molecular function | | Plasma membrane (ER/ extracellular) | |  | Endoplasmic reticulum (GO:0005783)  Chloroplast (GO:0009507) | Molecular_function_unknown (GO:0003674) | | | | Biological_process_unknown (GO:0008150) | |
| AT5G42146 | Unknown protein with unknown molecular function | | Plasma membrane | |  | Cellular_component_unknown (GO:0005575)  Mitochondrion (GO:0005739) | Molecular_function_unknown (GO:0003674) | | | | Biological_process_unknown (GO:0008150) | |
| AT3G29034 | Unknown protein with unknown molecular function | | Cytosol | |  | Cellular_component_unknown (GO:0005575)  Mitochondrion (GO:0005739) | Molecular_function_unknown (GO:0003674) | | | | Biological_process_unknown (GO:0008150) | |
| AT2G02180 | Tobamovirus multiplication protein 3 (TOM3): Unknown molecular function but necessary for the efficient multiplicatio  n of tobamoviruses | | Plasma membrane | |  | Mitochondrion (GO:0005739) |  | | | | Viral replication complex formation and maintenance (GO:0046786) | |
| **AGI** | **Description** | | **SUBAcon** | | **Ref** | **GO terms** | | | | | | |
|  |  |  |  |  |  | **Cellular component** | | **Molecular function** | | | | **Biological process** |
| **Unknown function continued** | | | | | | | | | | | | |
| AT1G25275 | Unknown hypothetical protein with unknown molecular function but is involved in karrikin response | | Extracellular | |  | Extracellular region (GO:0005576) | Molecular_function_unknown (GO:0003674) | | | | Response to karrikin (GO:0080167)  Regulation of defense response (GO:0031347)  Systemic acquired resistance (GO:0009627) | |
| AT4G02920 | Unknown protein with unknown molecular function | | Cytosol | |  | Mitochondrion (GO:0005739) | Molecular_function_unknown (GO:0003674) | | | | Biological_process_unknown (GO:0008150) | |
| **Potential false positives due to their localization** | | | | | | | | | | | | |
| AT5G52580 | RabGAP/TBC domain-containing protein involved in RAB GTPase activator activity | | Nucleus | |  | Cytoplasm (GO:0005737)  Cytosol (GO:0005829)  Intracellular (GO:0005622) | RAB GTPase activator activity (GO:0005097) | | | | Regulation of Rab GTPase activity (GO:0032313) | |
| AT1G52740 | Histone H2A protein 9 (HTA9): Involved in DNA methylation of transposons but not that of genes | | Nucleus | |  | Nucleosome (GO:0000786)  Nucleus (GO:0005634)  Vacuole (GO:0005773) | DNA binding (GO:0003677)  Protein binding (GO:0005515) | | | | Chromatin remodeling (GO:0006338)  Nucleosome assembly (GO:0006334)  Defense response to bacterium (GO:0042742)  Response to temperature stimulus (GO:0009266)  Regulation of flower development (GO:0009909) | |
| AT5G21170 | SNF1-related protein kinase regulatory subunit beta-1 (AKINBETA1): Encode a subunit of the SnRK1 kinase (Sucrose non-fermenting-1-related protein kinase). Involved in regulation of nitrogen and sugar metabolism. | | Nucleus | |  | Cytoplasm (GO:0005737) | Protein binding (GO:0005515)  AMP-activated protein kinase activity (GO:0004679) | | | | Response to sucrose stimulus (GO:0009744)  Response to fructose stimulus (GO:0009750)  N-terminal protein myristoylation (GO:0006499)  Cellular response to nitrogen levels (GO:0043562) | |
| AT3G10770 | Single-stranded nucleic acid binding R3H protein with unknown function | | Nucleus | |  | Cytoplasm (GO:0005737) | Nucleic acid binding (GO:0003676) | | | | Biological_process_unknown (GO:0008150) | |
| AT1G14980 | Chaperonin 10 (CPN10): Involved in protein folding in response to stress | | Mitochondria | |  | Mitochondrion (GO:0005739)  Cytoplasm (GO:0005737) | Copper ion binding (GO:0005507)  Chaperone binding (GO:0051087)  ATP binding (GO:0005524) | | | | Protein folding (GO:0006457)  Response to endoplasmic reticulum stress (GO:0034976)  Response to hydrogen peroxide (GO:0042542)  Response to heat (GO:0009408)  Response to high light intensity (GO:0009644) | |
| **AGI** | **Description** | | **SUBAcon** | | **Ref** | **GO terms** | | | | | | |
|  |  |  |  |  |  | **Cellular component** | | **Molecular function** | | | | **Biological process** |
| **Potential false positives due to their localization continued** | | | | | | | | | | | | |
| AT5G41700 | Ubiquitin-conjugating enzyme E2 8 (UBC8): Encodes one of the polypeptides that constitute the ubiquitin-conjugating enzyme E2 that’s involved in ubiquitination reaction to target protein to degradation. | | Peroxisome or cytosol or nucleus | |  | Nucleus (GO:0005634)  Cytoplasm (GO:0005737) | Ubiquitin-protein ligase activity (GO:0004842)  Acid-amino acid ligase activity (GO:0016881)  Protein binding (GO:0005515) | | | | Endosperm development (GO:0009960)  Ubiquitin-dependent protein catabolic process (GO:0006511)  Regulation of unidimensional cell growth (GO:0051510)  Proteasome assembly (GO:0043248)  Stamen development (GO:0048443)  Regulation of transport (GO:0051049)  DNA endoreduplication (GO:0042023)  Response to misfolded protein (GO:0051788)  Proteasomal ubiquitindependent protein catabolic process (GO:0043161) | |
| AT3G14290 | Proteasome subunit alpha type-5-B (PAE2): Encodes 20S proteasome subunit PAE2 (PAE2) that has RNase activity. | | Cytosol or nucleus | |  | Cytosol (GO:0005829)  Proteasome core complex (GO:0005839)  Cytosolic ribosome (GO:0022626)  Proteasome core complex, alpha-subunit complex (GO:0019773)  Cytoplasm (GO:0005737) | Ribonuclease activity (GO:0004540)  Threoninetype endopeptidase activity (GO:0004298)  Peptidase activity (GO:0008233) | | | | Ubiquitin-dependent protein catabolic process (GO:0006511)  Photorespiration (GO:0009853)  Fatty acid beta-oxidation (GO:0006635)  Proteolysis involved in cellular protein catabolic process (GO:0051603) | |
| AT4G37830 | Cytochrome c oxidase subunit VIa: Protein with cytochrome-c oxidase activity | | Mitochondria | |  | Mitochondrion (GO:0005739)  Mitochondrial inner membrane (GO:0005743)  Mitochondrial respiratory chain complex IV (GO:0005751) | Cytochrome-c oxidase activity (GO:0004129) | | | |  | |
| AT3G47070 | Uncharacterized protein with unknown molecular function | | Plastid | |  | Chloroplast  (GO:0009507)  Thylakoid (GO:0009579)  Chloroplast thylakoid (GO:0009534)  Chloroplast thylakoid membrane (GO:0009535)  Chloroplast envelope (GO:0009941) |  | | | | Photosynthesis (GO:0015979)  Thylakoid membrane organization (GO:0010027)  Response to blue light (GO:0009637)  Chloroplast relocation (GO:0009902)  Divalent metal ion transport (GO:0070838)  Cellular cation homeostasis (GO:0030003)  ncRNA metabolic process (GO:0034660)  rRNA processing (GO:0006364)  Response to red light (GO:0010114) | |
| AT1G79040† | Encodes for the 10 kDa PsbR subunit of photosystem II (PSII) | | Plastid | |  | Chloroplast (GO:0009507)  Thylakoid (GO:0009579)  Chloroplast thylakoid (GO:0009534)  Thylakoid membrane (GO:0042651)  Chloroplast thylakoid membrane (GO:0009535)  Photosystem II (GO:0009523)  Oxygen evolving complex (GO:0009654) | Protein binding (GO:0005515) | | | | Photosynthesis (GO:0015979)  Photosystem II oxygen evolving complex assembly (GO:0010270)  Response to red light (GO:0010114)  Response to far red light (GO:0010218)  Response to blue light (GO:0009637)  Response to sucrose stimulus (GO:0009744)  Response to high light intensity (GO:0009644)  Regulation of proton transport (GO:0010155)  Cysteine biosynthetic process (GO:0019344) | |
| **AGI** | **Description** | | **SUBAcon** | | **Ref** | **GO terms** | | | | | | |
|  |  |  |  |  |  | **Cellular component** | | **Molecular function** | | | | **Biological process** |
| **Potential false positives due to their localization continued** | | | | | | | | | | | | |
| AT5G38420 | RUBISCO SMALL SUBUNIT 2B (RBCS2B): Encodes subunit of Rubisco | | Plastid | |  | Chloroplast (GO:0009507)  Chloroplast stroma (GO:0009570)  Chloroplast envelope (GO:0009941)  Chloroplast ribulose bisphosphate carboxylase complex (GO:0009573)  Thylakoid (GO:0009579)  Apoplast (GO:0048046)  Membrane (GO:0016020)  Cytosolic ribosome (GO:0022626) | Ribulose-bisphosphate carboxylase activity (GO:0016984) | | | | Response to blue light (GO:0009637)  Response to red light (GO:0010114)  Response to far red light (GO:0010218)  Carbon fixation (GO:0015977) | |
| AT5G17170 | Enhancer of sos3-1 (ENH1): Involved in electron carrier activity and metal ion binding | | Plastid | |  | Chloroplast (GO:0009507)  Chloroplast envelope (GO:0009941)  Chloroplast thylakoid (GO:0009534)  Chloroplast thylakoid membrane (GO:0009535) | Metal ion binding (GO:0046872) | | | | Photosystem II assembly (GO:0010207)  Response to blue light (GO:0009637)  Response to red light (GO:0010114)  Response to far red light (GO:0010218)  rRNA processing (GO:0006364)  Pentose-phosphate shunt (GO:0006098)  Plastid organization (GO:0009657) | |
| AT5G21430 | CHLORORESPIRATORY REDUCTION L (CRRL): A chaperone DnaJ-domain superfamily protein involved in heat shock protein binding | | Plastid | |  | Chloroplast (GO:0009507)  Chloroplast thylakoid membrane (GO:0009535) | Heat shock protein binding (GO:0031072) | | | |  | |
| AT3G63410† | ALBINO OR PALE GREEN MUTANT 1 (APG1): Encodes a MPBQ/MSBQ methyltransferase involved in methylation step of PQ biosynthesis. | | Plastid | | [9] | Plastid  (GO:0009536)  Chloroplast (GO:0009507)  Chloroplast envelope (GO:0009941)  Chloroplast inner membrane (GO:0009706) | Methyltransferase activity (GO:0008168)  S-adenosylmethionine-dependent methyltransferase activity (GO:0008757)  2-methyl-6-phytyl-1,4-benzoquinone methyltransferase activity (GO:0051741) | | | | Metabolic process (GO:0008152)  Isopentenyl diphosphate biosynthetic process, mevalonate-independent pathway (GO:0019288)  Glucosinolate biosynthetic process (GO:0019761)  Plastoquinone biosynthetic process (GO:0010236)  Vitamin E biosynthetic process (GO:0010189)  Phosphatidylglycerol biosynthetic process (GO:0006655) | |
| AT5G38410† | RUBISCO SMALL SUBUNIT 3B (RBCS3B): Encodes a subunit of the Rubisco | | Plastid | | [9] | Chloroplast (GO:0009507)  Chloroplast envelope (GO:0009941)  Chloroplast stroma (GO:0009570)  Chloroplast ribulose bisphosphate carboxylase complex (GO:0009573)  Thylakoid (GO:0009579)  Cytosolic ribosome (GO:0022626)  Cell wall (GO:0005618)  Apoplast (GO:0048046)  Membrane (GO:0016020) | Ribulosebisphosphate carboxylase activity (GO:0016984) | | | | Photosynthesis (GO:0015979)  Response to blue light (GO:0009637)  Response to red light (GO:0010114)  Response to far red light (GO:0010218)  Chloroplast ribulose bisphosphate carboxylase complex biogenesis (GO:0080158)  Carbon fixation (GO:0015977) | |
| **AGI** | **Description** | | **SUBAcon** | | **Ref** | **GO terms** | | | | | | |
|  |  |  |  |  |  | **Cellular component** | | **Molecular function** | | | | **Biological process** |
| **Potential false positives due to their localization continued** | | | | | | | | | | | | |
| AT1G67740 | PsbY precursor (processed into two integral membrane proteins with identical topology, PsbY-1 and PsbY-2) and is involved in manganese ion binding | | Plastid | |  | Photosystem II (GO:0009523)  Chloroplast (GO:0009507)  Chloroplast photosystem II (GO:0030095)  Chloroplast thylakoid membrane (GO:0009535)  Chloroplast stromal thylakoid (GO:0009533)  Integral to membrane (GO:0016021) | Manganese ion binding (GO:0030145) | | | | Photosynthesis (GO:0015979)  Photosystem II assembly (GO:0010207)  Photosynthesis, light reaction (GO:0019684)  Plastid organization (GO:0009657)  Regulation of protein dephosphorylation (GO:0035304)  rRNA processing (GO:0006364)  Cysteine biosynthetic process (GO:0019344) | |
| AT2G26500† | Putative cytochrome b6f complex subunit (petM): Protein with plastoquinol-plastocyanin reductase activity | | Plastid | | [9] | Chloroplast (GO:0009507)  Chloroplast thylakoid membrane (GO:0009535)  Cytochrome b6f complex (GO:0009512) | Plastoquinol-plastocyanin reductase activity (GO:0009496) | | | | Plastid organization (GO:0009657)  Photosystem II assembly (GO:0010207)  rRNA processing (GO:0006364) | |
| AT1G61520† | Light-harvesting complex I chlorophyll a/b binding protein 3 (LHCA3): A component of the main light harvesting chlorophyll a/b-protein complex of Photosystem II | | Plastid | | [9] | Chloroplast (GO:0009507)  Chloroplast thylakoid (GO:0009534)  Chloroplast thylakoid membrane (GO:0009535)  Thylakoid (GO:0009579)  Membrane (GO:0016020)  Plastoglobule (GO:0010287)  Light-harvesting complex (GO:0030076) | Chlorophyll binding (GO:0016168) | | | | Photosynthesis (GO:0015979)  Photosynthesis, light harvesting (GO:0009765)  Cysteine biosynthetic process (GO:0019344) | |
| AT5G54270† | Light-harvesting chlorophyll B-binding protein 3 (LHCB3): A component of the main light harvesting chlorophyll a/b-protein complex of Photosystem II | | Plastid | | [9] | Chloroplast (GO:0009507)  Chloroplast thylakoid membrane (GO:0009535)  Thylakoid (GO:0009579)  Chloroplast thylakoid (GO:0009534)  Membrane (GO:0016020)  Light-harvesting complex (GO:0030076) | Structural molecule activity (GO:0005198) | | | | Photosynthesis (GO:0015979)  Photosynthesis, light harvesting (GO:0009765) | |
| AT4G21770 | RNA pseudourine synthase 6: Involved in RNA modification | | Plastid | |  | Chloroplast (GO:0009507) | Pseudouridine synthase activity (GO:0009982)  RNA binding (GO:0003723) | | | | Pseudouridine synthesis (GO:0001522)  RNA modification (GO:0009451) | |
| AT3G21055† | Photosystem II subunit T (PSBTN): Encodes photosystem II 5 kD protein subunit PSII-T | | Plastid | | [9] | Chloroplast (GO:0009507)  Chloroplast thylakoid lumen (GO:0009543)  Chloroplast photosystem II (GO:0030095) | Molecular_function_unknown (GO:0003674) | | | | Biological_process_unknown (GO:0008150)  rRNA processing (GO:0006364)  Plastid organization (GO:0009657)  Photosystem II assembly (GO:0010207)  Photosynthesis (GO:0015979)  Regulation of protein dephosphorylation (GO:0035304)  Photosynthesis, light reaction (GO:0019684) | |
| **AGI** | **Description** | | **SUBAcon** | | **Ref** | **GO terms** | | | | | | |
|  |  |  |  |  |  | **Cellular component** | | **Molecular function** | | | | **Biological process** |
| **Potential false positives due to their localization continued** | | | | | | | | | | | | |
| AT4G13590 | Uncharacterized protein with unknown molecular function | | Plastid | |  | Chloroplast (GO:0009507)  Chloroplast envelope (GO:0009941)  Chloroplast inner membrane (GO:0009706)  Membrane (GO:0016020) |  | | | |  | |
| AT5G53560 | Encodes a cytochrome b5 isoform that can be reduced by CBR, a cytochrome b5 reductase | | ER (PM, vacuole, Plastid) | |  | Endoplasmic reticulum (GO:0005783)  Endoplasmic reticulum membrane (GO:0005789)  Plasma membrane (GO:0005886)  Vacuole (GO:0005773)  Vacuolar membrane (GO:0005774)  Chloroplast thylakoid membrane (GO:0009535) | Heme binding (GO:0020037) | | | | Pentacyclic triterpenoid biosynthetic process (GO:0019745)  Sterol biosynthetic process (GO:0016126) | |
| AT2G29650 | Sodium-dependent phosphate transport protein 1 (PHT4;1): Encodes an inorganic phosphate transporter | | Plastid | |  | Chloroplast (GO:0009507)  Chloroplast thylakoid membrane (GO:0009535)  Thylakoid (GO:0009579)  Membrane (GO:0016020) | Sugar:hydrogen symporter activity (GO:0005351)  Inorganic diphosphate transmembrane transporter activity (GO:0030504)  Carbohydrate transmembrane transporter activity (GO:0015144)  Inorganic phosphate transmembrane transporter activity (GO:0005315)  Organic anion transmembrane transporter activity (GO:0008514) | | | | Nitrate transport (GO:0015706)  Defense response to bacterium (GO:0042742)  Jasmonic acid mediated signaling pathway (GO:0009867)  Transmembrane transport (GO:0055085)  Detection of biotic stimulus (GO:0009595)  Protein targeting to membrane (GO:0006612)  Response to chitin (GO:0010200)  Response to light stimulus (GO:0009416)  Defense response to fungus (GO:0050832)  Regulation of plant-type hypersensitive response (GO:0010363)  Systemic acquired resistance, salicylic acid mediated signaling pathway (GO:0009862)  Salicylic acid biosynthetic process (GO:0009697)  Regulation of hydrogen peroxide metabolic process (GO:0010310)  Negative regulation of defense response (GO:0031348)  Regulation of response to biotic stimulus (GO:0002831)  Response to nematode (GO:0009624)  MAPK cascade (GO:0000165)  Cellular response to water deprivation (GO:0042631)  Regulation of multiorganism process(GO:0043900) | |
| AT3G12610 | DNA-DAMAGE REPAIR/TOLERATION 100 (DRT100): Plays role in DNA-damage repair/toleration and can partially complements RecA- phenotypes | | Extracellular (plastid) | |  | plasma membrane  (GO:0005886)  chloroplast  (GO:0009507) | nucleotide binding  (GO:0000166) | | | | response to drug (GO:0042493)  signal transduction (GO:0007165)  response to chemical stimulus (GO:0042221)  response to UV (GO:0009411)  UV protection (GO:0009650) | |
| **AGI** | **Description** | | **SUBAcon** | | **Ref** | **GO terms** | | | | | | |
|  |  |  |  |  |  | **Cellular component** | | **Molecular function** | | | | **Biological process** |
| **Potential false positives due to their localization continued** | | | | | | | | | | | | |
| AT3G61470† | Photosystem I light harvesting complex protein (LHCA2) Encodes a component of the light harvesting antenna complex of photosystem I | | Plastid | | [9] | Chloroplast (GO:0009507)  Thylakoid (GO:0009579)  Chloroplast thylakoid (GO:0009534)  Chloroplast thylakoid membrane (GO:0009535)  Membrane (GO:0016020)  Lightharvesting complex (GO:0030076)  Photosystem I antenna complex (GO:0009782) | Chlorophyll binding (GO:0016168) | | | | Photosynthesis (GO:0015979)  Photosynthesis, light harvesting (GO:0009765)  Photosynthesis, light harvesting in photosystem I (GO:0009768) | |
| AT1G20340† | DNA-DAMAGE-REPAIR/TOLERATION PROTEIN 112 (DRT112): Plastocyanin protein involved in recombination and DNA-damage resistance | | Plastid | | [9] | Chloroplast (GO:0009507)  Thylakoid (GO:0009579)  Chloroplast thylakoid (GO:0009534)  Thylakoid lumen (GO:0031977)  Chloroplast thylakoid lumen (GO:0009543)  Chloroplast stroma (GO:0009570) | Electron carrier activity (GO:0009055)  Copper ion binding (GO:0005507) | | | | Positive regulation of transcription, DNAdependent (GO:0045893)  Response to UV (GO:0009411)  Cell differentiation (GO:0030154)  Hydrogen peroxide catabolic process (GO:0042744)  Anthocyanin accumulation in tissues in response to UV light (GO:0043481)  Response to copper ion (GO:0046688)  Regulation of translation (GO:0006417)  Cysteine biosynthetic process (GO:0019344)  Leaf morphogenesis (GO:0009965)  Response to blue light (GO:0009637)  Response to red light (GO:0010114)  Response to far red light (GO:0010218)  Multidimensional cell growth (GO:0009825)  Response to cadmium ion (GO:0046686)  Regulation of hormone levels (GO:0010817)  Pentose-phosphate shunt (GO:0006098)  Cell tip growth (GO:0009932)  Copper ion homeostasis (GO:0055070)  Regulation of proton transport (GO:0010155)  Response to high light intensity (GO:0009644)  Plastid organization (GO:0009657)  Response to chemical stimulus (GO:0042221)  Cell wall organization (GO:0071555)  Negative regulation of translation (GO:0017148)  Root hair elongation (GO:0048767)  Response to sucrose stimulus (GO:0009744)  Polysaccharide biosynthetic process (GO:0000271) | |
| Brackets in location row indicate alternative location consensus in SUBAcon [10,11]; † - identified as sticky proteins | | | | | | | | | | | | |
